# A novel cell-based sensor detecting the activity of individual basic proprotein convertases

**DOI:** 10.1101/464032

**Authors:** Karin Löw, Kornelia Hardes, Chiara Fedeli, Nabil G. Seidah, Daniel B. Constam, Antonella Pasquato, Torsten Steinmetzer, Alexandre Roulin, Stefan Kunz

## Abstract

The basic proprotein convertases (PCs) furin, PC1/3, PC2, PC5/6, PACE4, PC4, and PC7 are promising drug targets for human diseases. However, developing selective inhibitors remains challenging due to overlapping substrate recognition motifs and limited structural information. Classical drug screening approaches for basic PC inhibitors involve homogeneous biochemical assays using soluble recombinant enzymes combined with fluorogenic substrate peptides and do not accurately recapitulate the complex cellular context of the basic PC-substrate interaction. We report here *PCific*, a novel cell-based molecular sensor that allows rapid screening of candidate inhibitors and their selectivity toward individual basic PCs within mammalian cells. *PCific* consists of *Gaussia* luciferase linked to a sortilin-1 membrane anchor via a cleavage motif that allows efficient release of luciferase specifically if individual basic PCs are provided in *cis*. Screening of selected candidate peptidomimetic inhibitors revealed that *PCific* can readily distinguish between general and selective PC inhibitors in a high-throughput screening format.

## INTRODUCTION

The proprotein convertases (PCs) are a family of nine conserved calcium-dependent serine endoproteases that includes the basic PCs PC1/3, PC2, furin, PC4, PACE4, PC5/6, and PC7, as well as the non-basic SKI-1/S1P, and PCSK9 (Seidah & Prat, 2002, Seidah & Prat, 2007, Seidah & Prat, 2012). All PCs have a modular structure comprising a signal peptide, an N-terminal prodomain followed by a catalytic domain containing the “Ser/His/Asp” catalytic triad that catalyzes peptide bond scission, and a P domain. Furin, PC5/6B, PC7, and SKI-1/S1P have a transmembrane and C-terminal cytosolic domain. The soluble PC1/3 and PC2 accumulate in acidified secretory granules, and PC4, PC5/6A, PACE4, and PCSK9 are by default secreted (Seidah, 2011). Processing by PCs is essential for the proper function of many cellular proteins including pro-hormones, and precursors of growth factor, transcription factors, proteases, and adhesion molecules. Basic PCs have similar but not identical recognition motifs containing the minimal *consensus* sequence K/R-X_n_-K/R↓ and a large body of evidence indicates non-redundant roles *in vivo* (Seidah, Mayer et al., 2008, Seidah & Prat, 2012).

PCs are synthesized as inactive zymogens that require autoproteolytic processing for activation. After removal of the signal peptide in the ER, the prodomains of basic PCs are autoproteolytically cleaved, resulting in a latent form of the enzymes that remains associated with the inhibitory cleaved prodomains. Activation of the enzymes requires a second cleavage and release of the prodomain that occurs in specific sub-cellular locations (Anderson, Molloy et al., 2002, Creemers, Vey et al., 1995, Seidah et al., 2008). Crucial functions in development and normal physiology link PCs to a range of human disorders (Seidah & Prat, 2012). Many cellular substrates of basic PCs are involved in cell proliferation, malignant transformation, and metastasis formation, linking their expression to cancer progression and invasion (Artenstein & Opal, 2011, Jaaks & Bernasconi, 2017). Basic PCs can directly or indirectly contribute to atherosclerosis (Stawowy & Fleck, 2005), neurodegenerative diseases (Bennett, Denis et al., 2000, Lopez-Perez, Seidah et al., 1999), and to the regulation of inflammation (Wu, Song et al., 2016). Moreover, several human pathogens hijack basic PCs as essential cellular factors required for their multiplication (Stieneke-Grober, Vey et al., 1992, Thomas, 2002).

Considering the potential of basic PCs as drug targets, the development of specific inhibitors is of high priority. The first basic PC inhibitors were irreversible substrate-analogue chloromethyl ketones (CMK), in particular decanoyl-RVKR-CMK with high potency and preferred selectivity for basic PCs (Hallenberger, Bosch et al., 1992). Macromolecular inhibitor designs included insertion of basic PC K/R-Xn-K/R↓ motifs into the reactive site loop of α1-antitrypsin (Fugere & Day, 2005) or eglin c (Komiyama & Fuller, 2000). Screening of peptide-based inhibitors from libraries yielded potent polyarginines that inhibit basic PCs at low nanomolar concentrations (Cameron, Appel et al., 2000). More recent peptidomimetic designs resulted in several potent reversible competitive substrate analogue inhibitors derived from the lead structure phenylacetyl-Arg-Val-Arg-4-amidinobenzylamide that inhibit basic PCs at low nanomolar and even picomolar concentrations (Becker, Hardes et al., 2011, Becker, Lu et al., 2012, Becker, Sielaff et al., 2010, Hardes, Becker et al., 2015).

Despite a few multibasic 2,5-dideoxystreptamine derivatives with low nanomolar inhibition constants (Jiao, Cregar et al., 2006), most currently available non-peptide small molecule inhibitors for basic PCs show only moderate potency and cross-react between basic PCs, restricting therapeutic applications (Couture, Kwiatkowska et al., 2015). Conventional biochemical approaches using chromogenic or fluorogenic substrates and purified soluble PCs are suitable for high-throughput screening (HTS) and led to the discovery of several synthetic small molecule inhibitors. However, these HTS assays do not accurately recapitulate the complex cellular context of the basic PC-substrate interaction. Indeed, a more recent molecular sensor that can quantify PC activities in different sub-cellular compartments revealed that the local sub-cellular environment and spatial compartmentalization of basic PCs critically influences their activities and specificities (Bessonnard et al., 2015; Ginefra et al., 2018; Mesnard and Constam, 2010; Mesnard et al., 2011). This Cell-Linked Indicator of Proteolysis (CLIP) combines defined sub-cellular trafficking with monitoring of fluorescence resonance energy transfer (FRET) between suitable pairs of fluorophores in high-resolution imaging (Ginefra, Filippi et al., 2018). The prototypic CLIP consists of secreted enhanced cyan fluorescence protein (eCFP) and mCitrine fused *via* a flexible linker containing a canonical basic PC recognition motif (Mesnard & Constam, 2010) and allows detection of autocrine and paracrine PC activities at the plasma membrane (Mesnard, Donnison et al., 2011). Recently published CLIP designs containing cellular targeting signals allow quantitative assessment of basic PC bioactivities within specific subcellular compartments (Ginefra et al., 2018). Recent studies with these sensors uncovered complementary subcellular distribution of furin and PC7 bioactivities (Ginefra et al., 2018). In order to inhibit the activity of basic PC in the cellular context, inhibitors must reach critical concentrations within the subcellular compartment(s) containing the bioactive enzyme. Within a defined subcellular compartment, the local milieu, defined by pH, redox potential, and ion concentrations, may further influence inhibitor activity. Moreover, the format of classical HTS assay restricts screening to compounds that inhibit the catalytic activity of the mature enzyme. A cell-based assay opens a broader target-range for potential candidate inhibitors, including biosynthesis and zymogen activation, as well as cellular regulatory factors of basic PCs. The development of cell-based HTS platforms appears therefore promising. Here, we report *PCific*, a novel cell-based molecular sensor that allows rapid screening of collections of inhibitors and their selectivity toward individual basic PCs within mammalian cells.

## RESULTS

### Design and subcellular distribution of a latent biosensor of basic PCs

Previously, we developed a cell-based biosensor of the non-basic PC SKI-1/S1P that included an N-terminal *Gaussia* luciferase (GLuc) reporter fused to a SKI-1/S1P-derived membrane anchor *via* a cleavable peptide sequence mimicking a physiological SKI-1/S1P site (da Palma, Burri et al., 2014). Functional tests revealed that the sensor recapitulates key features of authentic substrates, including strict SKI-1/S1P specificity and correct subcellular location of processing (da Palma et al., 2014, da Palma, Cendron et al., 2016, Oppliger, da Palma et al., 2015). In the present study, we used this design as starting point to develop a cell-based basic PC sensor. However, human cell lines commonly used for high throughput small molecule screening co-express several members of the basic PC family at varying combinations and levels (Bessonnard, Mesnard et al., 2015). A major challenge for the development of a cell-based basic PC sensor was therefore the ability to discriminate between individual basic PCs present in the same cell. To overcome this problem, we sought to design a cell-based sensor containing a basic PC recognition sequence that is only inefficiently cleaved by endogenous basic PCs. Specific cleavage of the sensor will be achieved by overexpression of individual PCs in the same cell (in *cis*), followed by detection of a sensitive reporter released into the supernatant.

Most basic PCs process their substrates within the secretory pathway or at the cell surface at neutral or mildly acidic pH. Notable exceptions are PC1/3 and PC2 which are active within acidified secretory granules at an optimum pH <5.5 (Seidah & Prat, 2012). A major substrate of PC1/3 and PC2 in neuroendocrine tissues is the prohormone proopiomelanocortin (POMC). Specific cleavage of POMC by PC1/3 and PC2 at multiple basic PC recognition sites yields distinct peptides, including α, β, and γ-melanocyte stimulating hormone (Cawley, Li et al., 2016) (Fig. 1A). We hypothesized that the acidic pH requirement of PC1/3 and PC2 may render at least some POMC-derived cleavage sites relatively resistant to other endogenous basic PCs. To address this issue, we produced human POMC in HEK293T cells that express endogenous PC2, furin, PACE4, PC5/6, and PC7 (Bessonnard et al., 2015). Despite the absence of secretory granules in this cell type (Seidah & Chretien, 1999), POMC underwent efficient processing at most cleavage sites, with the notable exception of site 3 that flanks γ3-MSH and is processed by PC1/3 (Fig. 1B). As expected, co-expression of POMC with recombinant PC1/3 and PC2 resulted in weak, but significant processing at site 3 (Fig. 1B). The inefficient processing of the POMC site 3-derived recognition sequence by endogenous PCs made it a possible candidate for a “latent” cleavage site. To test this, we fused a GLuc reporter to a peptide containing POMC residues 91-121 (P14-P17’) flanking the POMC cleavage site 3 (Fig. 1C). The putative N-glycosylation motif (NSS) at position P14 may further restrict processing by endogenous PCs by glycan shielding (Bachert & Linstedt, 2013, Schjoldager, Vester-Christensen et al., 2011). The membrane-anchored basic PCs differ in their subcellular distribution (Ginefra et al., 2018, Seidah & Prat, 2012). Our sensor required therefore a broadly specific targeting sequence, ideally congruent with the individual basic PCs, to allow maximum co-localization with different family members. For this purpose, we chose a membrane anchor derived from sortilin-1. This protein traffics through most secretory compartments, reaches the cell surface, and undergoes endocytosis, followed by delivery to endosomes (Morinville, Martin et al., 2004, Willnow, Petersen et al., 2008). To minimize “off-target” cleavage by other proteases, we retained only the transmembrane domain and cytosolic tail (residues 756-831) of sortilin-1, necessary and sufficient for correct subcellular targeting (Fig. 1C). For the sake of simplicity, we name our sensor henceforth *PCific*, an acronym formed by the terms *PC* and *specific*. In a first step, we examined the sub-cellular distribution of the sensor. To this end, HeLa cells were transiently transfected with *PCific* bearing a V5 peptide-tag, fixed after 48 h, permeabilized, and examined by immunofluorescence. To assess subcellular localization of *PCific*, we performed double-stains combining anti-V5 antibody to the sensor with a panel of specific antibodies to markers for individual sub-cellular compartments or marker constructs. Consistent with the known broad subcellular distribution of sortilin-1, *PCific* co-localized with markers of ER, Golgi, trans-Golgi network (TGN), and partially with early and late endosomes (Fig. 2).

**Figure 1.**
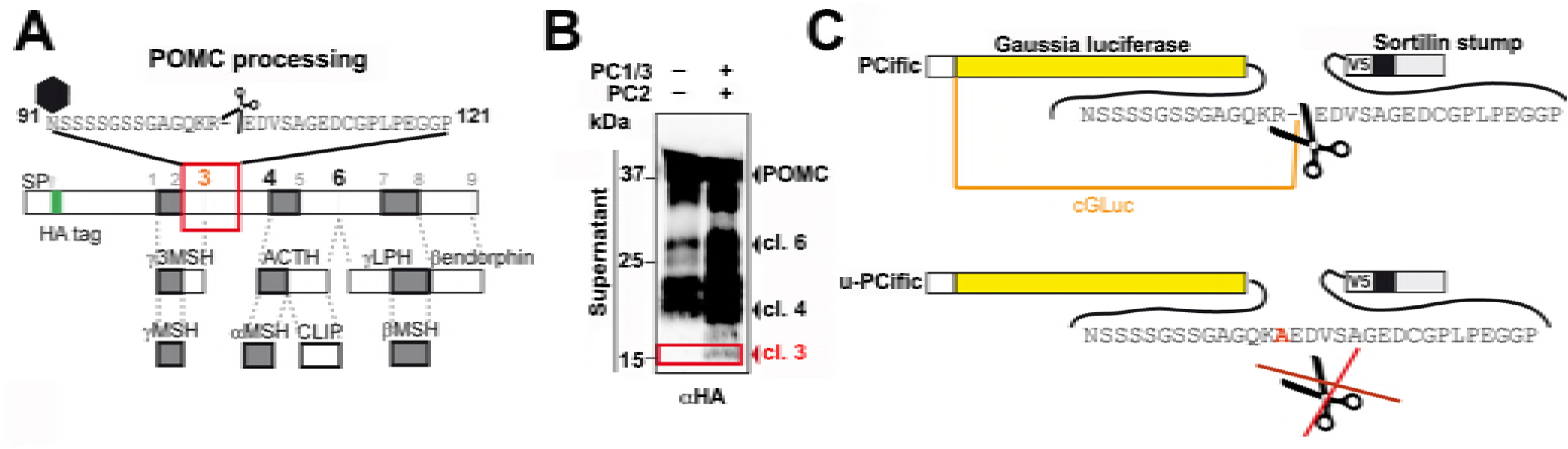
Design of *PCific*. (A) Schematic representation of POMC processing at cleavage sites 1-9 by PC1/3 and PC2. The cleavage products α-, β- and γ-MSH, ACTH, γ -LPH, and β- endorphin are indicated. The location of the cleavable sequence motif NSSSSGSSGAGQKR↓EDVSAGEDCGPLPEGGP present in *PCific* is outlined in red. The scissile bond (scissors) and the putative N-glycosylation site (black diamond) at P14 are indicated. (B) Inefficient processing of POMC cleavage site 3 by endogenous basic PCs. HEK293T cells were transfected with N-terminally HA-tagged human POMC in combination with either recombinant human PC1/3 and PC2, or with empty vector, and the cleavage pattern was assessed in Western blot after 72 h. The POMC precursor and the N-terminal cleavage fragments ending at sites 6, 4, and 3 are indicated. Note the apparent absence of the site 3 fragment in the lane lacking PC1/3 and PC2. (C) Schematic of *PCific* sensor (top) and its uncleavable version (below). The N-terminal GLuc is linked to the transmembrane and cytosolic domains of sortilin-1 via the POMC-derived sequence NSSSSGSSGAGQKR↓EDVSAGEDCGPLPEGGP, followed by a V5 peptide tag for detection. The R to A mutation in the dibasic cleavage motif of the uncleavable *PCific* variant (*u-PCific*) is highlighted in red.

**Figure 2.**
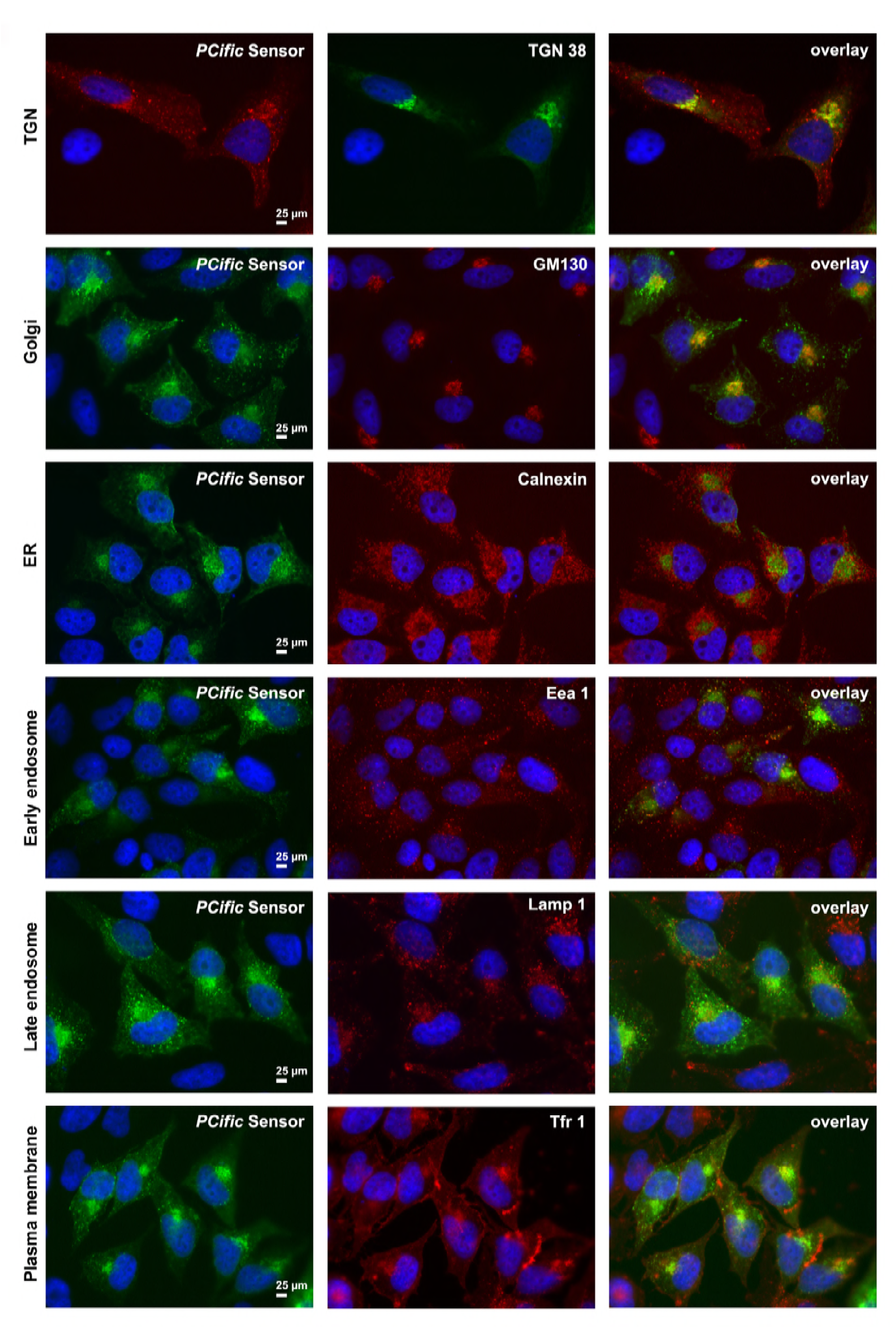
Subcellular localization of *PCific*. *PCific* was transiently expressed in HEK293T cells. After 48 h, cells were fixed, permeabilized, and *PCific* detected with a goat polyclonal antibody to the V5 epitope combined with Alexa Fluor 488 donkey anti-goat secondary antibody (green). The different subcellular compartments were labeled using specific primary antibodies against the indicated markers in combination with suitable Alexa Fluor 594- conjugated secondary antibodies (red) as detailed in Supplementary Materials and Methods. The trans-Golgi network (TGN) was labeled by expression of a fusion protein of citrine with the TMD and cytosolic tail of TGN 38. Here *PCific* sensor was detected with a mouse monoclonal antibody to the V5 epitope combined with Alexa Fluor 594 donkey anti-mouse secondary antibody (red). Nuclei were stained with NucBlue^TM^, bar = 25 μM.

### The latent basic PC sensor is specifically cleaved by furin and PC7 provided in *cis*

Furin represents the prototypic basic PC and is currently the most important drug target (Seidah & Prat, 2012). Furin undergoes activation in the TGN and can recycle between endosomes and TGN, involving sorting motifs in the cytosolic domain, recognized by specific adaptor proteins (Anderson et al., 2002, Chia, Gasnereau et al., 2011, Creemers et al., 1995, Mallet & Maxfield, 1999, Molloy, Thomas et al., 1998, Molloy, Thomas et al., 1994, Seidah et al., 2008, Takahashi, Nakagawa et al., 1995, Wan, Molloy et al., 1998). Co-transfection of *PCific* with recombinant furin in HEK293T and HeLa cells resulted in largely overlapping subcellular distribution (Fig. 3A). A possible concern was that over-expression of recombinant PCs may affect subcellular location. We therefore validated the location of recombinant furin and found extensive co-localization with markers of Golgi and TGN, as well as partial overlaps with early and late endosomes (Fig. S1), in line with published reports (Molloy et al., 1994, Takahashi et al., 1995, Wan et al., 1998). In sum, our co-localization studies confirmed the expected broad cellular distribution of *PCific* that overlaps with furin and likely other basic PCs.

**Figure 3.**
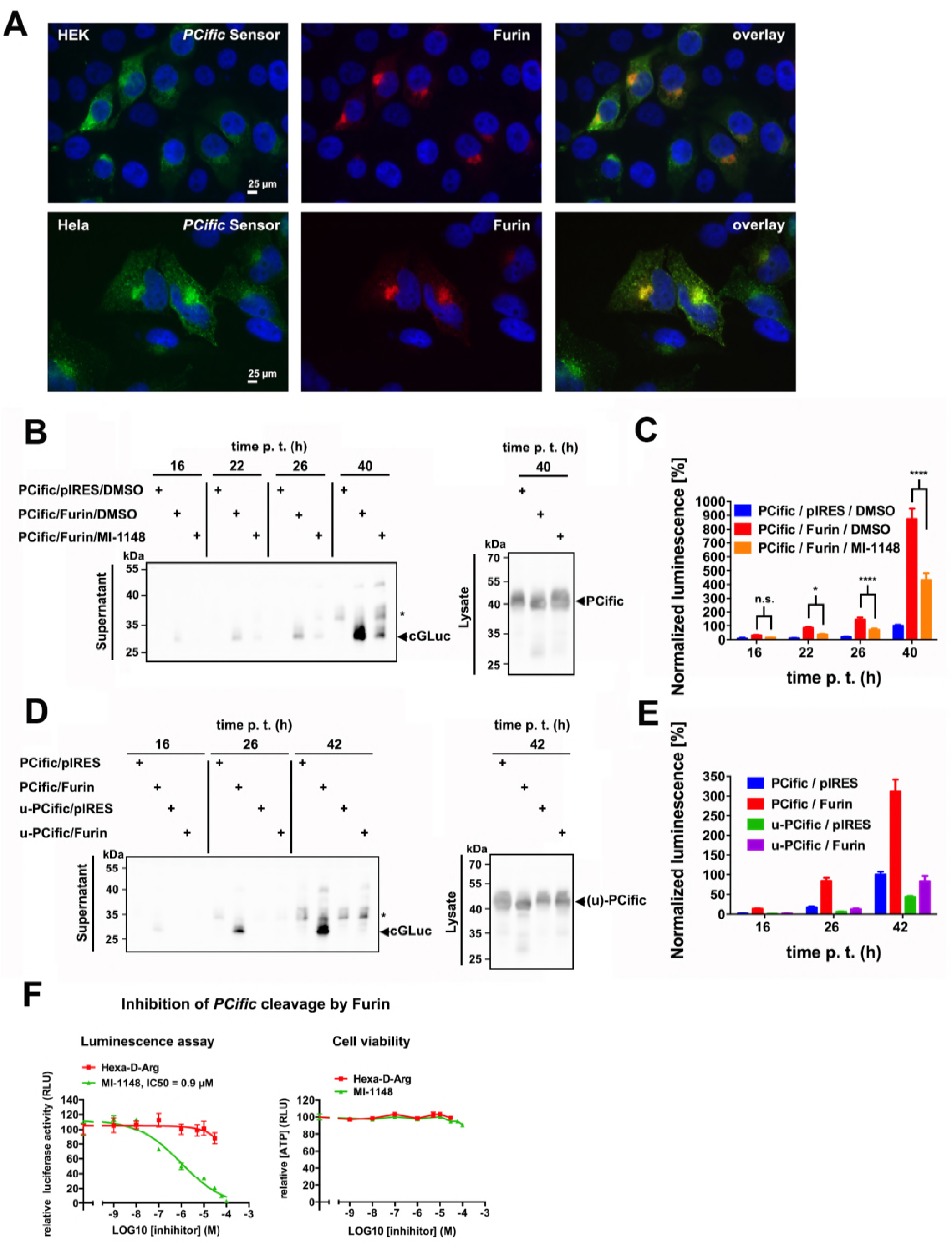
Specific cleavage of *PCific* by furin provided in *cis*. (A) Co-localization of *PCific* with recombinant furin. HEK293T and HeLa cells were co-transfected with *PCific* and recombinant human furin. After 48 h cells were fixed, permeabilized, and stained with a specific rabbit antibody to furin combined with Alexa Fluor 594-conjugated secondary antibodies (red). *PCific* (green) was detected as in Figure 2, and nuclei stained with NucBlue^TM^, bar = 25 μM. Merged images show that the broad distribution of *PCific* comprises the furin compartment. (B) Specific cleavage of *PCific* by furin provided in *cis*. HEK293T cells were transfected with *PCific* either in combination with furin or empty vector (pIRES). The basic PC inhibitor MI-1148 (25 μM) or DMSO solvent was added at 4 h post transfection. Cell culture supernatant was sampled at the indicated time points and cells lysed after 40 h, followed by Western blot using anti-GLuc antibody. Uncleaved *PCific* was detected in cell lysates (ca. 40 kDa), whereas cleaved GLuc (cGLuc) was found in supernatants (arrowhead, ca. 30 kDa). An unspecific GLuc positive species (*) was observed in the absence of furin or after furin inhibition. (C) Detection of GLuc activity in supernatants of (B) by luciferase assay. Data were recorded as arbitrary RLU and normalized to *PCific* transfected cells in the absence of furin and inhibitor (*PCific*/pIRES/DMSO) at 40 h; means ± SD; n = 9. (D) The uncleavable *PCific* variant (*u-PCific*) is resistant to furin provided in *cis*. HEK293T cells were transfected with *PCific* or *u-PCific*, combined with either furin or empty vector (pIRES). Supernatants harvested at the indicated time points and cell lysate harvested at 42 h were probed in Western blot as in (B). The cGLuc fragment and the “off-target” cleavage product (*) are indicated. Please note the absence of specific cleavage of *u-PCific*, even in presence of furin over-expression. (E) Detection of GLuc activity in supernatants of (D) by luciferase assay. Data were recorded as arbitrary RLU and normalized to *PCific* transfected cells in the absence of furin (*PCific*/pIRES) at 42 h; means ± SD; n = 9. (F) HEK293T cells were transfected with *PCific* sensor and furin plasmids and 0 - 100 μM of the inhibitors Hexa-D-arginine (Hexa-D-Arg) or MI-1148 were added 1 h post transfection. Supernatants were harvested 22 h p. t., luminescence from cGLuc activity was measured, corrected for background luminescence in the absence of furin and normalized to furin cleavage in the absence of inhibitor (n = 3, SEM). Dose response curves were established by non-linear regression curve fit and IC50 calculated from fit curves. Cell viability in presence of the different inhibitor concentrations was assessed via ATP levels in cell lysates corresponding to the supernatants in by CellTiter Glo^®^ assay (n = 3, SEM). No toxicity was observed for the doses tested.

Next, we verified the ability of our sensor to detect the bioactivity of furin in the cellular context. Briefly, HEK293T cells were transfected with *PCific*, either in combination with furin, or vector control. To verify furin-mediated processing, we used the recently described tight-binding inhibitor MI-1148 (Hardes et al., 2015). Cell supernatants were sampled at different time points post-transfection and release of GLuc monitored by Western-blot and luciferase assay. Detection of sensor expression levels in cell lysates revealed similar transfection efficiencies (Fig. 3B, right panel). In absence of recombinant furin, we observed basal levels of sensor processing that we operationally defined as background. Co-expression of furin induced the release of a cGLuc fragment of 30 kDa (Fig. 3B) and significant increase of luciferase activity in the supernatant over time (Fig. 3C). Treatment with the furin inhibitor MI-1148 (Hardes et al., 2015) reduced shedding of the cGLuc fragment and diminished the released luciferase activity (Fig. 3B, C). Next, the Arg residue in position P1 of our sensor was mutated to Ala to yield an uncleavable version (u-*PCific*) (Fig. 1C). The mutation reduced sensor cleavage by furin, assessed by Western blot (Fig. 3D) and luminescence assay (Fig. 3E). Expression of cleavable *PCific* in absence of exogenous furin or of u-*PCific* alone resulted in appearance of a GLuc positive species of higher molecular mass in the supernatant at late time points (Fig. 3B, D, asterisk), suggesting minor “off-target” processing. To define the dynamic range of our assay, HEK293T cells were co-transfected with *PCific* and furin, followed by treatment with increasing amounts of the furin inhibitor MI-1148 (Hardes et al., 2015) and hexa-D-arginine, that preferentially blocks basic PC activity in the extracellular space (Cameron et al., 2000). Cell viability was verified using CellTiter Glo^®^ assay that measures intracellular ATP levels as detailed in Materials and Methods. At the concentrations used, none of the inhibitors showed significant toxicity (Fig. 3F). Sensor processing was detected via released GLuc activity. The furin inhibitor MI-1148 almost completely reduced specific furin-dependent *PCific* cleavage in a dose-dependent manner with apparent IC50 value of 0.9 μM (Fig. 3F). In contrast, addition of up to 30 μM hexa-D-arginine had only a mild effect (Fig. 3F).

To further optimize assay conditions, we titrated the ratio of sensor plasmid relative to furin plasmid used for transfection and varied sampling time. Visualization of the results by a heat map revealed that improved signal-to-noise ratios were obtained at higher sensor/furin ratios and after short sampling times (Fig. 4A). To assess feasibility of HTS with our assay, robustness was determined *via* its Z’-factor (Z’= 1-(3σ^c+^+ 3σ^c-^)/(μ_c+_-μ_c-_), which depends on the sum of the standard deviations of positive and negative controls (σ_c+_ and σ_c-_, respectively) as well as the difference between the mean activity of these controls (μ_c+_ and μ_c-_). Assays with a Z’-factor ≥ 0.5 are considered “excellent” for HTS (Zhang, Chung et al., 1999). Using transient transfection of the sensor with furin as positive (μ_c+_) and empty vector as negative control (μ_c-_), and sampling of supernatants 22 h post transfection we obtained Z’ values between 0.63 (Fig. 3C) and 0.74 (Fig. 6A) for *PCific*, making our sensor highly suitable for HTS. In addition, we included PC7 that represents the phylogenetically most distant basic PC and an interesting drug target in its own right. Similar to furin, overexpression of PC7 resulted in enhanced processing of the cleavable sensor that was reduced by mutation of the cleavage site (Fig. 5A, B). At short sampling times, a higher sensor/PC7 ratio increased the signal-to-noise ratio (Fig. 5C), similar to the situation with furin (Fig. 4A). At later time points, a lower proportion of sensor *vs.* PC7 seemed favorable (Fig. 5C), suggesting a more complex situation when compared to furin. Under optimal assay conditions, PC7-specific cleavage of *PCific* likewise occurred with a Z’-factor of 0.74 (Fig. 7D), amenable for HTS.

**Figure 4.**
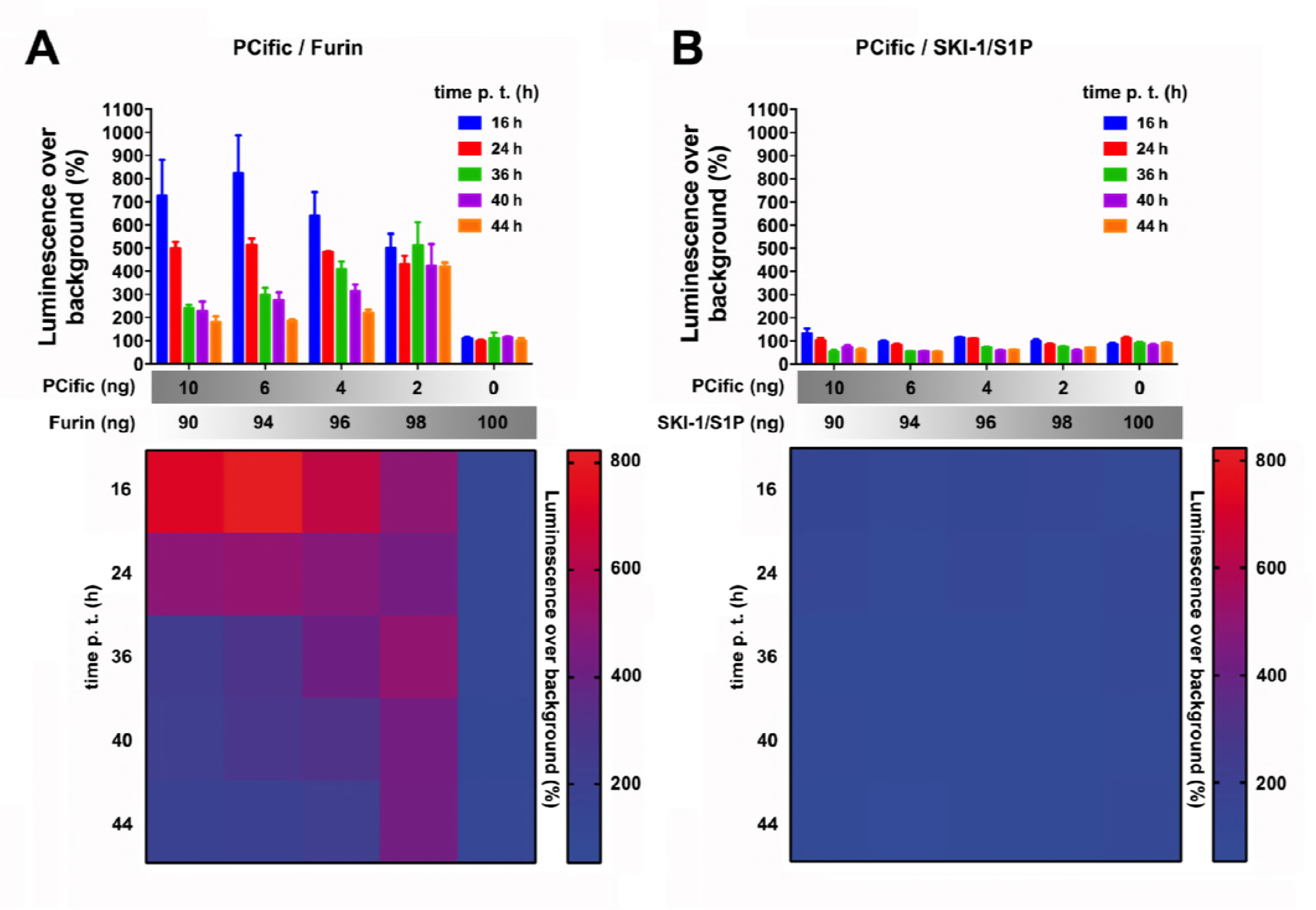
Optimization of conditions for furin-mediated *PCific* processing. HEK293T cells were transfected with the indicated ratios of *PCific* to furin plasmid (A) or *PCific* to SKI-1/S1P plasmid (B) as negative control. Background luminescence was assessed in cells transfected with the corresponding plasmid ratios of *PCific* to empty vector (pIRES). Cell supernatants were harvested at the indicated time points and luminescence measured as arbitrary RLU, followed by normalization to background luminescence, which was set at 100%. Experiments were carried out in biological triplicates with three independent measurements of luciferase activity. Mean values and SD are shown (n = 3). Data are depicted as heat maps below the corresponding bar diagrams with specific luminescence over background color coding. Sampling time (y-axis) is plotted against the sensor/PC plasmid ratio (x axis).

**Figure 5.**
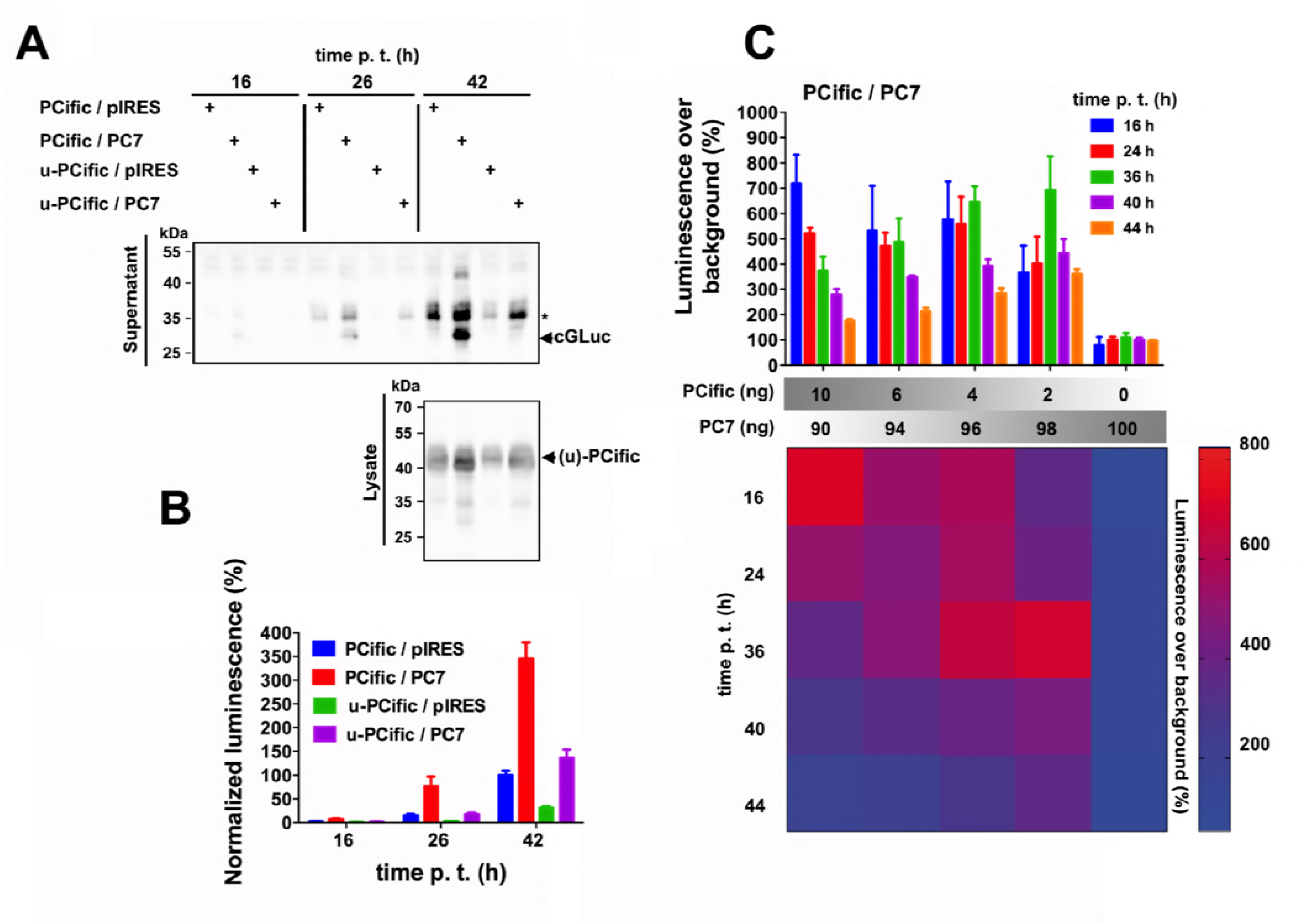
Cleavage of *PCific* by PC7 provided in *cis*. (A) HEK293T cells were transfected with *PCific* (cleavable or uncleavable version) either in combination with PC7 or empty vector (pIRES). Cell culture supernatant was sampled at the indicated time points and cells lysed after 42 h, followed by Western blot as in 3D. The specific cGLuc fragment (arrowhead) and the unspecific GLuc positive species are indicated (*). (B) Detection of GLuc activity in supernatants of (A) by luciferase assay. Data were recorded as RLU and normalized to *PCific* transfected cells in the absence of PC7 (*PCific*/pIRES) at 42 h; means ± SD; n = 9. (C) Identification of optimal conditions for PC7-mediated cleavage of *PCific.* Luminescence in cell supernatants was determined for different *PCific*/PC7 plasmid ratios at the indicated time points and expressed as fold-increase over background defined in *PCific/p*IRES transfected cells. Mean values ± SD, n = 3. Bottom: Results displayed as heat map as in 4A, B.

### Implementation of PCific in an inhibitor screen

The robust, rapid, and cost-effective assay format make the *PCific* sensor a promising candidate for a HTS platform to identify novel basic PC inhibitors from compound libraries. For proof-of-concept, we employed *PCific* to screen a selected set of peptidomimetic basic PC inhibitors that act as reversible substrate analogues (Hardes et al., 2015, Ivanova, Hardes et al., 2017) (Table 1). The lead structure of our compound library was the previously developed inhibitor phenylacetyl-Arg-Val-Arg-4-amidinobenzylamide MI-0227 (Becker et al., 2011, Becker et al., 2010). Modifications of the MI-0227 scaffold included incorporation of basic groups at position P5, introduction of the non-natural amino acid *tert*-leucine (Tle) in position 3, as well as introduction of less basic residues in position P1 (Becker et al., 2012, Hardes et al., 2015, Ivanova et al., 2017) (Table 1). In a forward screen, HEK293T cells were co-transfected with *PCific* and furin, followed by treatment with inhibitors at a concentration of 25 μM (Fig. 6A, B). Cell viability was monitored under the exact assay conditions by CellTiter Glo^®^ assay and overtly toxic compounds excluded (Fig. 6C). Most inhibitors significantly blocked furin-specific *PCific* processing in our HTS assay at the used concentration of 25 μM (Fig. 6B) with the notable exception of hexa-D-arginine, in line with our previous results (Fig. 3F). For counter-screening, we tested our candidates against *PCific* processing by PC7 and against our SKI-1/S1P sensor. As expected based on our previous studies, almost all candidates were inactive against the non-basic PC SKI-1/S1P that processes substrates at hydrophobic recognition sites (Seidah, Mowla et al., 1999) (Fig. 6G, H). The inhibition profile of the candidate compounds against PC7 was more complex and overlapped only partially with furin (Fig. 6D-F). Plotting the relative activities of the individual compounds against furin and PC7 revealed that the inhibitors used fell into three categories: 1) compounds with apparent selectivity for furin over PC7, 2) candidates with partial selectivity, and 3) compounds lacking detectable specificity (Fig. 6I). For validation, we chose two compounds of the first category, the parental compound MI-0227 (Fig. 6J) and compound MI-1530 (Fig. 6K). Performing dose-response characteristics, we confirmed that MI-0227 and MI-1530 show significant selectivity for furin over PC7, respectively.

**Figure 6.**
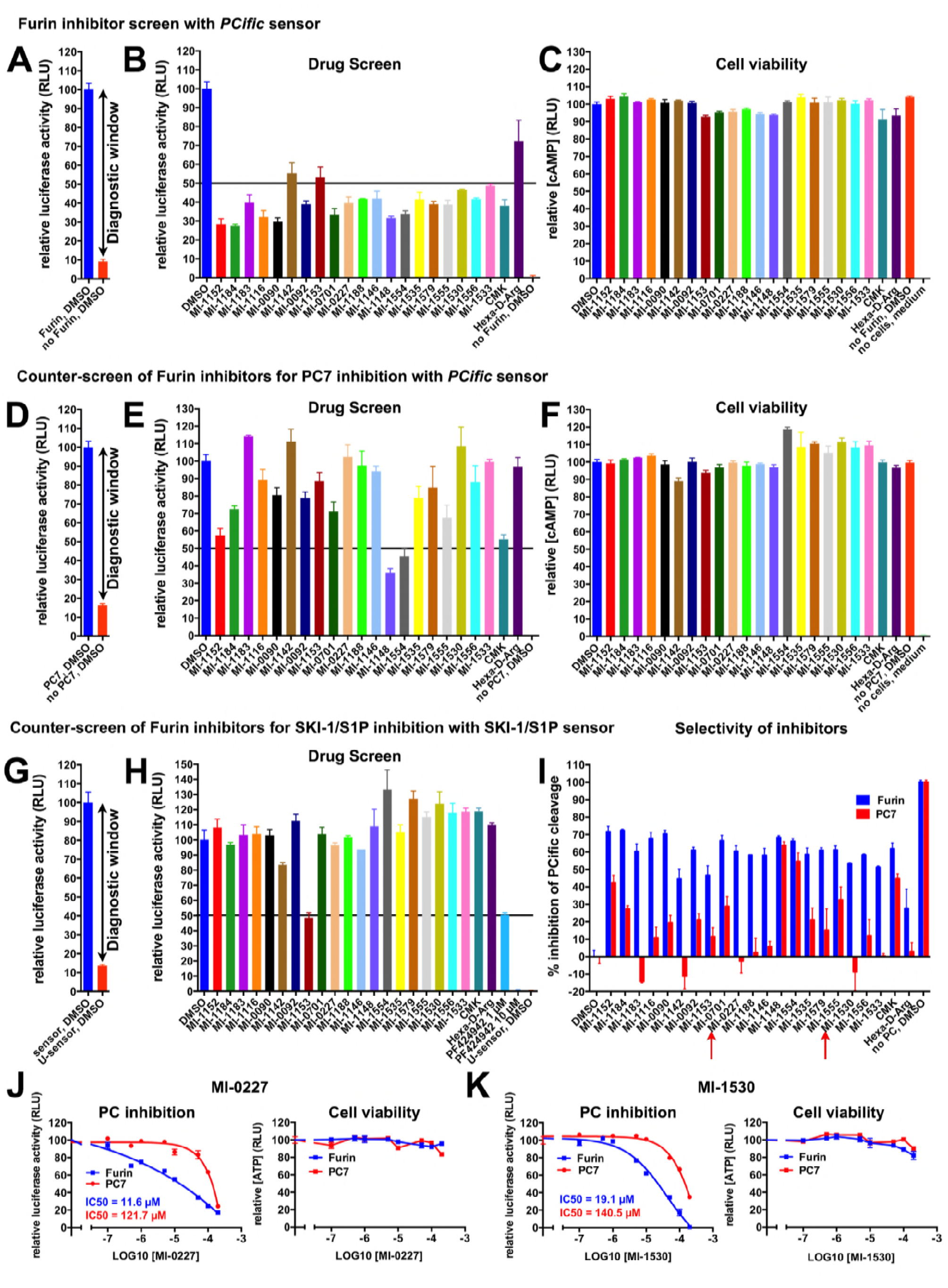
Differential inhibition of furin and PC7 cleavage of *PCific* by candidate compounds. (A-H) A selection of peptidomimetic compounds previously identified to inhibit fluorogenic peptide cleavage by recombinant soluble furin in *in vitro* assays (Marburg inhibitors (MI)) was subjected to *in vivo* screening for furin (A, B) and PC7 inhibition (D, E), using *PCific* sensor. Basic PC specificity was further confirmed in a counter-screen using SKI-1/S1P sensor (G, H). HEK293T cells were transfected with *PCific* and furin (A, B, C), *PCific* and PC7 (D, E, F), or SKI-1/S1P sensor plasmid (G, H) and inhibitor candidates were added 4 h p. t. to final concentrations of 25 μM, 0.25% DMSO. PC inhibition was analyzed by detection of cGLuc activity in cell supernatants by luminescence measurement (RLU) at 22 h post transfection. Diagnostic windows for the individual assays were established by comparison of the normalized RLU in supernatants of *PCific* and pIRES2-EGFP transfected cells to *PCific* and furin (A) or *PCific* and PC7 (B) transfected cells, both in absence of inhibitors (0.25% DMSO only). The diagnostic window for inhibition of endogenous SKI-1/S1P was determined by normalization of RLU in supernatants of cells transfected with non-cleavable SKI-1/S1P sensor to RLU obtained with cleavable SKI-1/S1P sensor, both in absence of inhibitors (G). For drug screening assays (B, E, H), luminescence values were background subtracted (red bars in A, D, G) and normalized to furin - (B), PC7-(E) and endogenous SKI-1/S1P cleavage (H) in the absence of inhibitor (0.25% DMSO) (n = 3, SEM), which was set to 100 %. Suppression of RLU to ≤ 50% was set as a threshold for efficient inhibition (black lines in B, E, H). Toxicity of the tested inhibitors at 25 μM was investigated in cell viability assays using the corresponding cell lysates from furin and PC7 screening analyses as described in Fig. 3F (n = 3, SEM) (C, F). The selectivity of furin over PC7 inhibition was assessed by calculating the percentage of inhibition for each compound at 25 μM from results shown in B and E. Mean values for furin and PC7 inhibition are depicted side by side in a bar diagram for individual MI-compounds (n = 3, SEM) (I). Two inhibitors with high furin over PC7 selectivity at 25 μM, MI-0227 and MI-1530 (red arrows in I), were selected to establish dose response curves for furin and PC7 inhibition using cleavage of *PCific* sensor as a read-out. A concentration range of 0 - 200 μM was tested (n =3, SEM) (J, K). IC50 values were determined after non-linear regression curve fit, confirming furin selectivity of MI-0227 and MI-1530 with approximately 10 and 7 fold lower IC50 values for furin compared to PC7 inhibition, respectively (J, K). Potential toxicity of inhibitors at the applied doses was excluded by accompanying cell viability assays with the corresponding cell lysates (J, K).

**Table 1.** List of inhibitors used in the study. For details, please see text.

**Figure 6. Differential inhibition of furin and PC7 cleavage of PCific by candidate compounds.**
| Inhibitor | Sequence | K<sub>i</sub>-value (nM)<br>against furin | # in Publication | Publication |
| --- | --- | --- | --- | --- |
| MI-1152 | 5-Gva-Tle-Arg-4-amidinobenzylamide | 1.26 | 30 | Hardes et al., ChemMedChem 2017 |
| MI-1184 | H-dArg-Arg-Arg-Val-Arg-4-amidinobenzylamide | 0.0283 | 9 | Hardes et al., ChemMedChem 2017 |
| MI-1183 | 4-AMe-Phac-Arg-Glu-Arg-4-amidinobenzylamide | 2.68 |  | Not published |
| MI-1116 | Phac-Arg-Tle-Arg-4-amidinobenzylamide | 0.17 | 9 | Hardes et al., ChemMedChem 2015 |
| MI-0090 | 4-GMe-Phac-Arg-Tle-Arg-Noragmatin | 0.28 | 25 | Ivanova et al., ChemMedChem 2017 |
| MI-1142 | Ac-YGRKKRRQRRVR-4-amidinobenzylamide | 11.99<sup>1</sup> |  | Not published |
| MI-0092 | 4-GMe-Phac-Arg-Tle-Arg-Agmatin | 0.22 | 26 | Ivanova et al., ChemMedChem 2017 |
| MI-1153 | nona-D-arginine (D9R) | 1.3 |  | Kacprzak et al., JBC 2004 |
| MI-0701 4 | 4-GMe-Phac-Arg-Val-Arg-4-amidinobenzylamide | 0.0076 | 17 | Becker et al., JBC 2012 |
| MI-0227 3 | Phac-Arg-Val-Arg-4-amidinobenzylamide | 0.66 | 3 | Hardes et al., ChemMedChem 2015<br>Becker et al., JMedChem 2010 |
| MI-1188 3 | 4-GMe-Phac-Arg-Glu-Arg-4-amidinobenzylamide | 0.58 |  | Not published |
| MI-1146 | 5-Gva-Val-Arg-4-amidinobenzylamide | 2.54 | 28 | Hardes et al., ChemMedChem 2017 |
| MI-1148 4 | 4-GMe-Phac-Arg-Tle-Arg-4-amidinobenzylamide | 0.0055 | 19 | Hardes et al., ChemMedChem 2015 |
| MI-1554 4 | 4-GMe-Phac-Arg-Tle-Lys-4-amidinobenzylamide | 0.0085 | 27 | Ivanova et al., ChemMedChem 2017 |
| MI-1535 | 4-GMe-Phac-Arg-Val-Arg-4-amidomethyl-2-aminopyridin | 8.32 | 16 | Ivanova et al., ChemMedChem 2017 |
| MI-1579 | 4-GMe-Phac-Arg-Tle-Arg-(3-fluor)-4-amidinobenzylamide | 0.0223 | 24 | Ivanova et al., ChemMedChem 2017 |
| MI-1555 | 4-GMe-Phac-Arg-Tle-Arg-5-amidomethyl-2-aminopyridin | 0.80 | 20 | Ivanova et al., ChemMedChem 2017 |
| MI-1530 | 4-GMe-Phac-Arg-Val-Arg-p-diaminoxilen | 3.47 | 13 | Ivanova et al., ChemMedChem 2017 |
| MI-1556 | 4-GMe-Phac-Arg-Tle-Arg-p-diaminoxilen | 2.52 | 19 | Ivanova et al., ChemMedChem 2017 |
| MI-1533 | 4-GMe-Phac-Arg-Val-Arg-5-amidomethyl-2-aminopyridin | 3.73 | 15 | Ivanova et al., ChemMedChem 2017 |
| CMK | Dec-Arg-Val-Lys-Arg-CMK |  |  | Garten et al., 1994 Biochimie |
| D6R | hexa-D-arginine | 106<br>265 |  | Cameron et al., JBC 2000<br>Fugere et al., MolPharmacol 2007 |

<sup>1</sup> Binding mode unknown, therefore only an IC<sub>50</sub> value in nM could be determined.
CMK – chloromethylketone, Dec – decanoyl, GMe – guanidinomethyl, Gva – 5-guanidinovaleryl, Phac – phenylacetyl, Tle – *tert*-leucine

Based on the structural relationships between compounds analyzed (Table 1), we tried to assess structure-activity patterns related to furin/PC7 selectivity. For the original lead compound MI-0227, the observed selectivity for furin over PC7 in our assay was in line with the previously established selectivity profile assessed via an *in vitro* enzymatic assay (Becker et al., 2011, Becker et al., 2012, Becker et al., 2010). Modifications of the MI-0227 scaffold had significant effects on the furin/PC7 selectivity of compounds used in our cell-based assay (Table 2). The most striking effect was observed with the replacement of Val with Tle in position P3 that reduced furin/PC7 selectivity in all candidate compounds tested: MI-0227/MI-1116, MI-0701/MI-1148, MI-1530/MI-1556, MI-1533/MI-1555, MI-1146/MI-1152 (Fig. 6I and Table 2). Interestingly, replacement of Val with Tle in position P3 enhances potency against furin (Hardes et al., 2015), suggesting that the nature of the residue in position P3 may improve recognition of PC7 in the cellular context. Introduction of a positively charged para-guanidinomethyl substitution on the phenylacetyl residue in position P5 of MI-0227 enhanced potency for furin by >100-fold (Hardes et al., 2015), but seemed to reduce selectivity in our system (Fig. 6I and Table 2). Replacement of Val by Glu in position P3, which introduced a negative charge, e.g. in compounds MI-1188 and MI-1183, restored furin/PC7 selectivity, due to a significant reduction of PC7 inhibition, suggesting a role of the overall charge of the molecule. This seems in line with the observation that introduction of less basic groups in position P1 of MI-1148 in compounds MI-1556 and MI-1555, and of MI-0701 in compounds MI-1530 and MI-1533 (Ivanova et al., 2017) or removal of the positively charged Arg in position P4 (MI-1146) likewise restored furin/PC7 selectivity (Fig. 6I and Table 2). Although limited in scope, our screen demonstrates that our *PCific* sensor can readily distinguish between general and selective PC inhibitors in a HTS format.

**Table 2.** Selectivity Furin/PC7 of selected inhibitors detected in the PCific-based screening assay. For details, please see text.

| Inhibitor | Sequence | K<sub>i</sub>-value (nM)<br>against furin | Selectivity<br>Furin/PC7 | Publication |
| --- | --- | --- | --- | --- |
| MI-1116 | Phac-Arg-Tle-Arg-4-amidinobenzylamide | 0.17 | + | Hardes et al., ChemMedChem 2015 |
| MI-0227 | Phac-Arg-Val-Arg-4-amidinobenzylamide | 0.66 | ++ | Becker et al., JMedChem 2010 |
| MI-0701 | 4-GMe-Phac-Arg-Val-Arg-4-amidinobenzylamide | 0.0076 | (+) | Becker et al., JBC 2012 |
| MI-1148 | 4-GMe-Phac-Arg-Tle-Arg-4-amidinobenzylamide | 0.0055 | - | Hardes et al., ChemMedChem 2015 |
| MI-1554 | 4-GMe-Phac-Arg-Tle-Lys-4-amidinobenzylamide | 0.0085 | - | Ivanova et al., ChemMedChem 2017 |
| MI-1579 | 4-GMe-Phac-Arg-Tle-Arg-(3F)-4-amidinobenzylamide | 0.0223 | (+) | Ivanova et al., ChemMedChem 2017 |
| MI-1188 | 4-GMe-Phac-Arg-Glu-Arg-4-amidinobenzylamide | 0.58 | ++ | Not published |
| MI-1183 | 4-AMe-Phac-Arg-Glu-Arg-4-amidinobenzylamide | 2.68 | ++ | Not published |
| MI-1530 | 4-GMe-Phac-Arg-Val-Arg-p-diaminoxilen | 3.47 | ++ | Ivanova et al., ChemMedChem 2017 |
| MI-1556 | 4-GMe-Phac-Arg-Tle-Arg-p-diaminoxilen | 2.52 | + | Ivanova et al., ChemMedChem 2017 |
| MI-1533 | 4-GMe-Phac-Arg-Val-Arg-5-amidomethyl-2-aminopyridin | 3.73 | ++ | Ivanova et al., ChemMedChem 2017 |
| MI-1555 | 4-GMe-Phac-Arg-Tle-Arg-5-amidomethyl-2-aminopyridin | 0.80 | (+) | Ivanova et al., ChemMedChem 2017 |
| MI-1535 | 4-GMe-Phac-Arg-Val-Arg-4-amidomethyl-2-aminopyridin | 8.32 | (+) | Ivanova et al., ChemMedChem 2017 |
| MI-1146 | 5-Gva-Val-Arg-4-amidinobenzylamide | 2.54 | ++ | Hardes et al., ChemMedChem 2017 |
| MI-1152 | 5-Gva-Tle-Arg-4-amidinobenzylamide | 1.26 | (+) | Hardes et al., ChemMedChem 2017 |
| MI-0090 | 4-GMe-Phac-Arg-Tle-Arg-Noragmatin | 0.28 | + | Ivanova et al., ChemMedChem 2017 |
| MI-0092 | 4-GMe-Phac-Arg-Tle-Arg-Agmatin | 0.22 | (+) | Ivanova et al., ChemMedChem 2017 |
| MI-1184 | H-dArg-Arg-Arg-Val-Arg-4-amidinobenzylamide | 0.0283 | (+) | Hardes et al., ChemMedChem 2017 |
| CMK | Dec-Arg-Val-Lys-Arg-CMK |  | - | Garten et al., 1994 Biochimie |

CMK – chloromethylketone, Dec – decanoyl, GMe – guanidinomethyl, Gva – 5-guanidinovaleryl, Phac – phenylacetyl, Tle – *tert*-leucine

## DISCUSSION

Basic PCs are linked to a wide range of human pathologies, including metabolic disorders, cancer, and infectious diseases, making them attractive targets for therapeutic intervention (Seidah & Prat, 2012). The development of specific inhibitors is therefore of high priority. Screens of large combinatorial multibasic peptide libraries yielded candidate inhibitors selective for PC1 *vs*. PC2 or furin, proving the feasibility to develop relatively specific inhibitors against basic PCs (Apletalina, Appel et al., 1998, Cameron et al., 2000). Some selectivity against the different basis PCs was recently found for a series of reversible competitive peptidomimetic substrate analogue derivatives (Becker et al., 2011, Becker et al., 2012, Becker et al., 2010, Hardes et al., 2015, Ivanova et al., 2017). Whereas furin, PC1/3, PC4, PACE4 and PC5/6 were inhibited with similar strong potency a reduced affinity has been observed for PC7 and PC2. Combined with high-resolution structure analysis, these inhibitors provided first insights into the structural basis for the observed strong furin potency (Dahms, Arciniega et al., 2016, Dahms, Hardes et al., 2014, Dahms, Jiao et al., 2017, Hardes et al., 2015). However, the development of non-peptide small molecule basic PC inhibitors remains challenging and to the best of our knowledge, none of the currently available small molecule basic PC inhibitors is completely specific for one family member.

Classical HTS for PC inhibitors mostly rely on homogeneous biochemical assays that do not accurately recapitulate the cellular context. Spatial subcellular compartmentalization of basic PCs clearly influences their specificities towards many substrates (Bessonnard et al., 2015, Ginefra et al., 2018, Mesnard & Constam, 2010, Mesnard et al., 2011) and development of cell-based assays appears promising. A prototypic cell-based basic PC assay involved an alkaline phosphatase (AP) reporter linked to a Golgi-anchor domain *via* a furin cleavage site (Coppola, Hamilton et al., 2007). It remains however unclear to what extent this sensor-type allows distinction between general and selective PC inhibitors. In contrast, our design deliberately prevents efficient processing by endogenous basic PCs present in many mammalian cell lines. Efficient cleavage of our latent sensor *PCific* is only achieved by over-expressing individual basic PCs in *cis*, which likely facilitates co-localization. A suitable latent cleavable sequence was derived from the PC1/3 site 3 of POMC that underwent only inefficient processing by endogenous basic PCs. Since basic PCs have complex, partially non-overlapping cellular trafficking patterns, we included targeting signals derived from sortilin-1 to achieve broad distribution of *PCific* covering most sub-cellular compartments (Morinville et al., 2004, Willnow et al., 2008). Notably, in contrast to the existing CLIP sensors, the sub-cellular specificity of *PCific* is defined by the distribution pattern of the exogenously provided basic PC. To evaluate our sensor, we tested the ability of *PCific* to specifically detect the bioactivities of furin and PC7 provided in *cis.* Optimization of assay conditions resulted in a favorable signal-to-noise ratio with Z’-value of 0.74 for both furin and PC7, which is considered “excellent” for HTS (Zhang et al., 1999).

For proof-of-concept, we employed *PCific* to screen a selected panel of peptidomimetic basic PC inhibitors derived from the lead compound phenylacetyl-Arg-Val-Arg-4-amidinobenzylamide (MI-0227) that exhibits selectivity for furin over PC7 assessed by *in vitro* assay (Becker et al., 2011, Becker et al., 2010). MI-0227 showed likewise >10-fold selectivity for furin vs. PC7 in our cell-based assay. A first examination of the structure-activity relationship of the MI-0227 derivatives tested in our assay revealed an unexpected role of the Val residue in P3 for the selectivity for furin vs. PC7. In all scaffolds tested, replacement of Val with Tle consistently reduced furin/PC7 selectivity. The reasons for this are currently unclear, but the data suggest that the nature of the residue in P3 may influence the recognition by PC7. In addition, for some inhibitors, we observed an apparent correlation between the charge of the molecule and its furin/PC7 selectivity. Introduction of a negative charge in position P3 and introduction of less basic groups in P1 apparently enhanced furin/PC7 selectivity. While the cellular distribution of recombinant furin was largely in line with published data, there is very recent evidence that an excess of PC7 can alter its cellular trafficking (Ginefra et al., 2018). To directly test the influence of expression levels, we currently develop inducible stable cell lines that allow fine-tuning of PC7/sensor ratios suitable for HTS. In summary, the robust and cost-effective assay format of *PCific* makes it particularly suited to identify novel specific small molecule inhibitors against basic PCs for therapeutic application in HTS. Its cell-based nature will allow screening for drug targets in addition to the catalytically active mature enzyme, including maturation, transport, and cellular factors that modulate the enzyme’s activity. This broadened “target range” will enhance the likelihood to isolate novel small molecule compounds that inhibit the enzymes in a direct or indirect manner and represents a conceptual advantage. Lastly, *PCific* is suitable to analyze the selectivity of inhibitor candidates toward individual basic PCs within mammalian cells.

## MATERIAL AND METHODS

### Cell culture and transfections

Human embryonic kidney cells (HEK293T) were maintained in DMEM supplemented with 10% (vol/vol) FBS, 100 U/ml penicillin and 0.1 mg/ml streptomycin. Transfections of HEK293T cells with sensor and proprotein convertase/IRES GFP plasmids were performed using Lipofectamine 3000 according to the manufacturer’s instructions. Transfection efficiencies were evaluated by detection of enhanced green fluorescent protein (EGFP).

### Detection of sensor cleavage by Western blot and luciferase assay

Conditioned cell supernatants harvested at the indicated time points were cleared by centrifugation for 5 min at 1,500 rpm. Cell layers were washed with cold PBS and lysed in CelLytic^TM^ M buffer (Sigma-Aldrich GmbH, Buchs, Switzerland) supplemented with Complete^TM^ Protease Inhibitor Cocktail (F. Hoffmann-La Roche Ltd., Basel, Switzerland) for 30 min at 4°C. Cell lysates were centrifuged for 5 min at 13,000 rpm. Conditioned media and cell lysates were mixed with 6 x reducing SDS-PAGE buffer and boiled for 5 min. Proteins were separated by SDS-PAGE and blotted onto nitrocellulose. Membranes were blocked with 3% (wt/vol) skim milk powder in PBS, 0.2% (vol/vol) Tween 20 (Sigma), and probed with rabbit anti-GLuc antibody (1:2000), followed by incubation with secondary HRP-conjugated goat/swine anti-rabbit antibodies (1:3000). Secreted GLuc activity was detected in the collected cell supernatants as described (da Palma et al., 2014) and detailed in Supplementary Information.

### Screening of furin inhibitors with *PCific* sensor

HEK293T cells were transfected in 96 well plates with 10 ng *PCific* plasmid per well in combination with either 90 ng of furin-, PC7- or empty pIRES2-EGFP plasmid using Lipofectamine 3000. Selected inhibitor canditates were added to the cell culture medium 4 h post transfection to a final concentration of 25 μM, 0.25% DMSO. All inhibitors were tested in triplicate in three independent transfections. Supernatants were harvested 22 h post transfection and 18 h after inhibitor addition. 10 μl of conditioned supernatant were pre-laid in white, half-volume 96-well plates (Costar) and 60 μl of coelenterazine substrate solution (16 ng/ml in PBS) were automatically injected to each well. Cleaved Gaussia luciferase (cGLuc) activity was analyzed immediately thereafter by luminescence measurement for 0.1 s as described. Luminescence values in supernatants from *PCific* and pIRES2-EGFP transfected cells (no inhibitor, 0.25% DMSO) were used to establish background cleavage, and the mean value obtained from three independent transfections was subtracted from furin / PC7 mediated *PCific* cleavage in all wells, with and without inhibitor. Background corrected cleavage in the presence of inhibitor was then normalized to background corrected cleavage of *PCific* by furin / PC7 in the absence of inhibitor (0.25% DMSO), which was set to 100%. Mean values (n =3) and SEM of background corrected, normalized luciferase activity in cell supernatants are depicted. All candidate compounds suppressing luciferase activity to ≤ 50% at 25 μM were considered effective inhibitors. The percentage of background subtracted, normalized residual luciferase activity in presence of 25 μM inhibitor was subtracted from 100% to obtain the percentage of inhibition, which was then depicted side by side for furin and PC7 for each of the inhibitor candidates (n = 3, SEM). In this way selectivity of an inhibitor for furin over PC7 could be assessed. Per definition, values of 0 % inhibition were obtained in cells expressing *PCific* and furin/PC7 in the absence of inhibitor, and 100% inhibition in the absence of *PCific* cleavage, when no furin or PC7 were co-expressed with *PCific* sensor.

To confirm their specificity for basic PCs, inhibitors were further counter-screened for inhibition of endogenous SKI-1/S1P activity. To this end, HEK293T cells were transfected in 96 well plates with 100 ng/well of SKI-1/S1P sensor plasmid, or an uncleavable version thereof, serving as a negative control. Background luminescence in cell supernatants was established with uncleavable SKI-1/S1P sensor (0.25% DMSO) and was subtracted from the luminescence obtained with cleavable SKI-1/S1P sensor, with or without inhibitors. Luciferase activity was then normalized to background corrected cleavage of SKI-1/S1P sensor in the absence of inhibitor (0.25% DMSO), which was set to 100 %. Mean values and SEM for the relative luciferase activities are depicted for the individual inhibitors at 25 μM (n = 3).

### Dose-response curves

HEK293T cells were transfected with 10 ng *PCific* plasmid per well in combination with 90 ng of furin / pIRES2-EGFP plasmid, and inhibitors were added 1 h post transfection. Supernatants were harvested 22 h post transfection, and luminescence was determined as described above. Luminescence values were background subtracted and normalized to furin cleavage in the absence of inhibitor and plotted against inhibitor concentration (mean + SEM, n =3). IC50 values were determined after non-linear regression curve fit (variable slope, four parameters).

## ACKNOWLEDGEMENTS

The authors thank Bruno G.H. Filippi for his critical reading of the manuscript and his useful comments. This research was supported by an Interdisciplinary Research Grant of the Faculty of Biology and Medicine of the University of Lausanne to A.R. and S.K., Novartis Foundation for Biomedical Research Grant 14B062 to A.R., Swiss National Science Foundation grants 310030_170108 to S.K., 31003A_173178 and 31003A_153467 to A.R. and 31003A_156452 to D.B.C.

